# SLB-msSIM: a spectral library-based multiplex segmented SIM platform for single-cell proteomic analysis

**DOI:** 10.1101/2024.10.22.618936

**Authors:** Lakmini Senavirathna, Cheng Ma, Van-An Duong, Hong-Yuan Tsai, Ru Chen, Sheng Pan

## Abstract

Mass spectrometry (MS)-based single-cell proteomics, while highly challenging, offers unique potential for a wide range of applications to interrogate cellular heterogeneity, trajectories, and phenotypes at a functional level. We report here the development of the spectral library-based multiplex segmented selected ion monitoring (SLB-msSIM) method, a conceptually unique approach with significantly enhanced sensitivity and robustness for single-cell analysis. The single-cell MS data is acquired by multiplex segmented selected ion monitoring (msSIM) technique, which sequentially applies multiple isolation cycles with the quadrupole using a wide isolation window in each cycle to accumulate and store precursor ions in the C-trap for a single scan in the Orbitrap. Proteomic identification is achieved through spectral matching using a well-defined spectral library. We applied the SLB-msSIM method to interrogate cellular heterogeneity in various pancreatic cancer cell lines, revealing common and distinct functional traits among PANC-1, MIA-PaCa2, AsPc-1, HPAF, and normal HPDE cells. Furthermore, for the first time, our novel data revealed the diverse cell trajectories of individual PANC-1 cells during the induction and reversal of epithelial-mesenchymal transition (EMT). Collectively, our results demonstrate that SLB-msSIM is a highly sensitive and robust platform, applicable to a wide range of instruments for single-cell proteomic studies.

## 1. INTRODUCTION

Mass spectrometry (MS)-based single-cell proteomics offers a distinctive approach for system-wide, unbiased profiling of proteomes in individual cells or cell clusters, enabling functional investigations with cellular resolution. Over the past few years, significant efforts have been made in single-cell mass spectrometric analysis. These developments include label-free approaches [1–7], as well as isobaric labeling/carrier proteome-based approaches, such as SCoPE-MS[8–10], iBASIL[11], and SCeptre[12]. The ion mobility technique has also been utilized to improve the sensitivity in single-cell proteomic analysis[13–16]. It is notable that while the utility of an isobaric labeled carrier proteome can boost the MS signal for single-cell analysis, challenges and limitations have been indicated in association with such approaches[9, 17]. More recently, single-cell proteomic works leveraged by high-frequency instruments with ion mobility have demonstrated superior sensitivity and resolution in the analysis of single-cell proteomes[7, 14–16]. These innovations have greatly improved the sensitivity and utility of single-cell proteomics.

In conventional bulk-scale shotgun proteomics, peptide and protein identification primarily relies on matching MS/MS spectral data with in-silico digested peptides from the target proteome through a database search. This approach is naturally extended to single-cell proteomics. However, single-cell analysis poses significant challenges due to the minute amount of extracted proteins from each cell, resulting in an extremely low signal-to-noise ratio and brief elution time for peptide detection. Consequently, there is poor peptide fragmentation and low-quality MS/MS spectra in both data-independent acquisition (DIA) and data-dependent acquisition (DDA) analyses. To address these challenges inherent to single-cell proteomics, current efforts predominantly focus on enhancing MS/MS signal sensitivity through employing high-end instruments like timsTOF or Orbitrap Astral, which offer high-frequency, highly sensitive mass analyzers, and ion mobility capabilities, to improve MS/MS signal quality in single-cell analysis. Given that the MS1 precursor signal of a peptide is significantly stronger than its MS/MS signals generated through gas-phase fragmentation reactions, a critical question arises: can MS1 signals be effectively used for peptide identification in single-cell proteomic analysis using a well-defined spectral library?

In our previous study, we demonstrated a proof-of-concept spectral library-based platform for single-cell proteomics, which differs conceptually from most current approaches[18]. To overcome the challenge of low MS/MS signal intensity inherent in single-cell analysis, this method leverages MS1 data to extract comprehensive peptide information, including monoisotopic mass, isotopic distribution, and retention time (hydrophobicity), for single-cell proteomic analysis. This information, which specifically defines the physicochemical characteristics of a peptide, is used to match with an empirically generated spectral library for peptide sequence identification.

Here, we report a critical technical advance in the spectral library-based single-cell proteomic platform to significantly enhance its sensitivity and robustness for single-cell analysis by employing the multiplex segmented Selected Ion Monitoring (msSIM) technique for precursor acquisition. The method sequentially applies multiple isolation cycles with the quadrupole using a wide isolation window in each cycle to accumulate and store precursor ions in the C-trap for a single scan in the Orbitrap. The msSIM method is employed to obtain single-cell data for proteomic identification through spectral library matching, significantly enhancing sensitivity, coverage, and robustness in single-cell analysis. With this improved mass spectrometric method, we applied the spectral library-based msSIM (SLB-msSIM) platform to investigate the single-cell heterogeneity of pancreatic cancer cell lines and sought to better understand the implication of cellular heterogeneity of individual cells in epithelial-mesenchymal transition (EMT), a crucial hallmark for determining metastasis states[19, 20].

## 2. EXPERIMENTAL SECTION

### Cell culture

Pancreatic cancer cells, including PANC-1, MIA-PaCa2, AsPC-1 and HPAF, were maintained in DMEM (ATCC, Manassas, VA) supplemented with 10% fetal bovine serum (FBS) and 1% penicillin-streptomycin. Normal human pancreatic ductal epithelial cells (HPDE-c7) were maintained in Keratinocyte serum-free medium supplemented with bovine pituitary extract and epidermal growth factor (Invitrogen, Carlsbad, CA). After reaching 70–80% confluence in culture flasks, the cells were detached using trypsin, washed twice with PBS, and then gathered for proteomic analysis.

### TGFβ1 treatment

0.5×10^6^ of PANC-1 cells were seeded in 60 mm cell culture dishes with serum containing DMEM medium. The culture medium was switched to the serum-free medium 24 hrs before the TGFβ1 treatment. The cells were exposed to 10 ng/mL of TGFβ1 (R&D systems, Minneapolis, MN) in serum-free medium for 3 days. Following the 3-day TGFβ1 treatment, the medium was changed to serum-free medium without TGFβ1, and the cells were cultured for an additional 4 days (totaling 7 days) to assess the reversal effect of EMT. Single-cell samples were collected on control (Con, without TGFβ1 treatment), day 3 (D3, EMT), and day 7 (D7, EMT reversal).

### Western blot

Cells were lysed with M-PER lysis buffer (Thermo Fisher Scientific, Waltham, MA) with 1% protease inhibitor (Sigma, St. Louis, MA). Equal amounts of protein were loaded into SDS-PAGE gels (Bio-Rad, Hercules, CA) and protein was transferred to the PVDF membrane using an iBlot 2 machine (Thermo Fisher Scientific). The membranes were immersed in 5% milk in phosphate-buffered saline + Tween 20 to block non-specific background staining. The EMT marker antibodies E-cadherin (1:1000), Snail (1:1000), and Actin (1:5000) were obtained from Cell Signaling (Danvers, MA). Membranes were incubated with HRP-conjugated secondary antibody (Jackson Immuno Research Laboratories, West Grove, PA) and detected using chemiluminescent reagents (Thermo Fisher).

### Single-cell and bulk sample preparation

The single-cell sample preparation was carried out as previously described using the DIRECT method[18]. Briefly, single cells were transferred directly to a glass injection vial with 3 µL PBS. Cells were subjected to freeze-thaw cycles for three times by placing the vials on dry ice and thawing briefly on ice. After each thaw cycle, cells were sonicated for 3 min in an ultrasonic cold-water bath. Samples were treated with dithiothreitol (DTT) and iodoacetamide (IDA) followed by adding ammonium bicarbonate to adjust the pH (∼ 7.8). Samples were digested with MS-grade trypsin (Thermo Fisher Scientific) at 37 ^0^C overnight. The digested samples were collected at the bottom of the vial by briefly centrifuging them and the reaction was stopped by adding 0.1% formic acid.

For bulk scale analysis, cell lysates were collected with M-PER lysis buffer (Thermo Fisher Scientific) containing 1% protease inhibitor (Sigma, St. Louis, MA). Lysates were treated with 10 mM DTT and incubated for 1hr at 50 ^0^C, followed by alkylation with 25 mM IDA for 30 min at room temperature. Tri-chloroacetic acid (TCA) precipitation was performed to precipitate the proteins. The protein precipitate was suspended in 50 mM ammonium bicarbonate and digested with trypsin (1:30) at 37 ^0^C overnight. Samples were acidified with 0.1% formic acid to stop the reaction. Digested peptides were purified with c18 columns (NEST group, Ipswich, MA).

### LC-MS/MS analysis

LC-MS/MS analysis was performed using a Q Exactive HF-X Orbitrap mass spectrometer interfaced with an UltiMate 3000 Binary RSLCnano HPLC System (Thermo Fisher Scientific). For bulk scale analysis, one microgram of a sample was loaded into a 5 mm trap column packed with 5 µM/100 Å C18 material (Thermo Fisher Scientific) using 98% buffer A (0.1% formic acid in water) and 2% buffer B (0.1% formic acid in acetonitrile) at a flow rate of 3 µL/min. The samples were separated in a self-packed C18 analytical column (75 µm X 25 cm) packed with 3 µm Reprosil-Pur C18-AQ material (Dr. Maisch, Germany) using a 90-minute linear gradient from 2% to 35% buffer B versus buffer A at a flow rate of 350 nL/min. Mass spectrometric analysis was performed using DDA mode. The survey scan was performed with 60,000 resolution with *m/z* 400 to 1600 with an AGC target of 3e6 and a max injection time of 50 msec. Monoisotopic masses were then selected for fragmentation for the 25 most abundant precursor ions within a dynamic exclusion range of 30 seconds. The minimum intensity threshold was set as 2.3e5. Fragmentation priority was given to the most intense ions. Precursor ions were isolated using the quadrupole with an isolation window of 1.6 *m/z*. Higher energy collisional dissociation (HCD) was applied with a normalized collision energy of 28%.

Mass spectrometric analyses for single-cell samples were conducted with the msSIM approach, which used the t-SIM acquisition mode provided by Xcalibur and employed a wide isolation window of 50 Th for multiplex isolation of precursor ions with the quadrupole. In total, the method applied a multiplexing degree of 17 for the MS1 scan range of *m/z* 400 to 1250. The midpoint of each segmented mass isolation window was used as the target mass for the inclusion list (e.g., 425 *m/z* was used as the target mass for the mass selection window of *m/z* 400-450). The precursor ions sequentially isolated in each cycle with the quadrupole were accumulated in the C-trap, where they are stored together before being finally detected in the Orbitrap in one single scan. The MS1 data was obtained at a resolution of 30,000, with an AGC target of 5e4 and a maximum injection time of 250 msec.

### Construction of spectral library

Protein and peptide identification in the bulk samples was used to construct the spectral library for single-cell analysis. The DDA data was searched against the UniProt human protein database (UniProt ID: UP000005640 (canonical), released June 2020; 20,394 protein entries) using the Comet algorithm[21] embedded in the Trans-Proteomic Pipeline[22]. Carbamidomethylation of cysteine was set as a fixed modification, and oxidation of methionine was set as a variable modification. The peptide identification was validated with a PeptideProphet[23] error rate of 0.01 and an E-value of 0.05. The peptides and proteins identified in the bulk samples were used to build the spectral library using Skyline[24] (version 22.2.0.312). To enhance the peptide matching for single-cell analysis, internal retention time (iRT) calibration was performed by generating an iRT calibrator using 11 peptides[25] (**Supplementary Table S1**).

### Spectral library-based single-cell data analysis

The raw msSIM data of the single-cell samples were imported to Skyline for peptide and protein identification by spectral matching. Accurate matching was achieved by precursor mass, isotopic distribution and retention time (hydrophobicity). The peptide and transition settings were set as previously described[18]. Peptides with an Isotope Dot Product less than 0.5 were excluded from the analysis. The abundance of each peptide was normalized to the total precursor intensity and presented as ion per million (IPM) using the following formula: Normalized Intensity (IPM) = Peptide Intensity/Total precursor intensity × 1,000,000. To remove the outliers of single-cell samples, the median absolute deviation (MAD) method was applied[18]. A dataset with the absolute deviation of the total precursor intensity greater than 2xMAD would be considered as an “outlier” and removed from the sample group for further analysis. Protein quantification was achieved by summation of the normalized intensities of the corresponding peptides. The peptides and proteins with an average intensity of less than 0.01 IPM were excluded from the analysis.

### Statistical and bioinformatic analysis

The t-SNE (t-distributed Stochastic Neighbor Embedding) plots were created to illustrate the similarities and differences among individual cells. t-SNE analyses were performed using Morpheus based on Pearson correlation. Unpaired two-tailed Student’s t-tests were conducted to compare the protein abundances between two groups of individual cells. Hallmark genes were identified using Gene Set Enrichment Analysis (GSEA, version 4.3.3)[26]. The cnetplots of the protein network were generated using the enrichplot package in R (version 4.4.0) with genes showing core enrichment in GSEA analysis.

## 3. RESULTS

### The SLB-msSIM method for enhanced proteomic identification in single-cell analysis

The SLB-msSIM workflow for single-cell analysis is illustrated in **Figure 1A**, consisting of the following modules: I) the “all-in-one” DIRECT method for single-cell preparation[18], II) a high-resolution spectral library generated from a bulk-scale DDA analysis, III) LC-msSIM data acquisition, and IV) spectral library-based data analysis.

**Figure 1.**
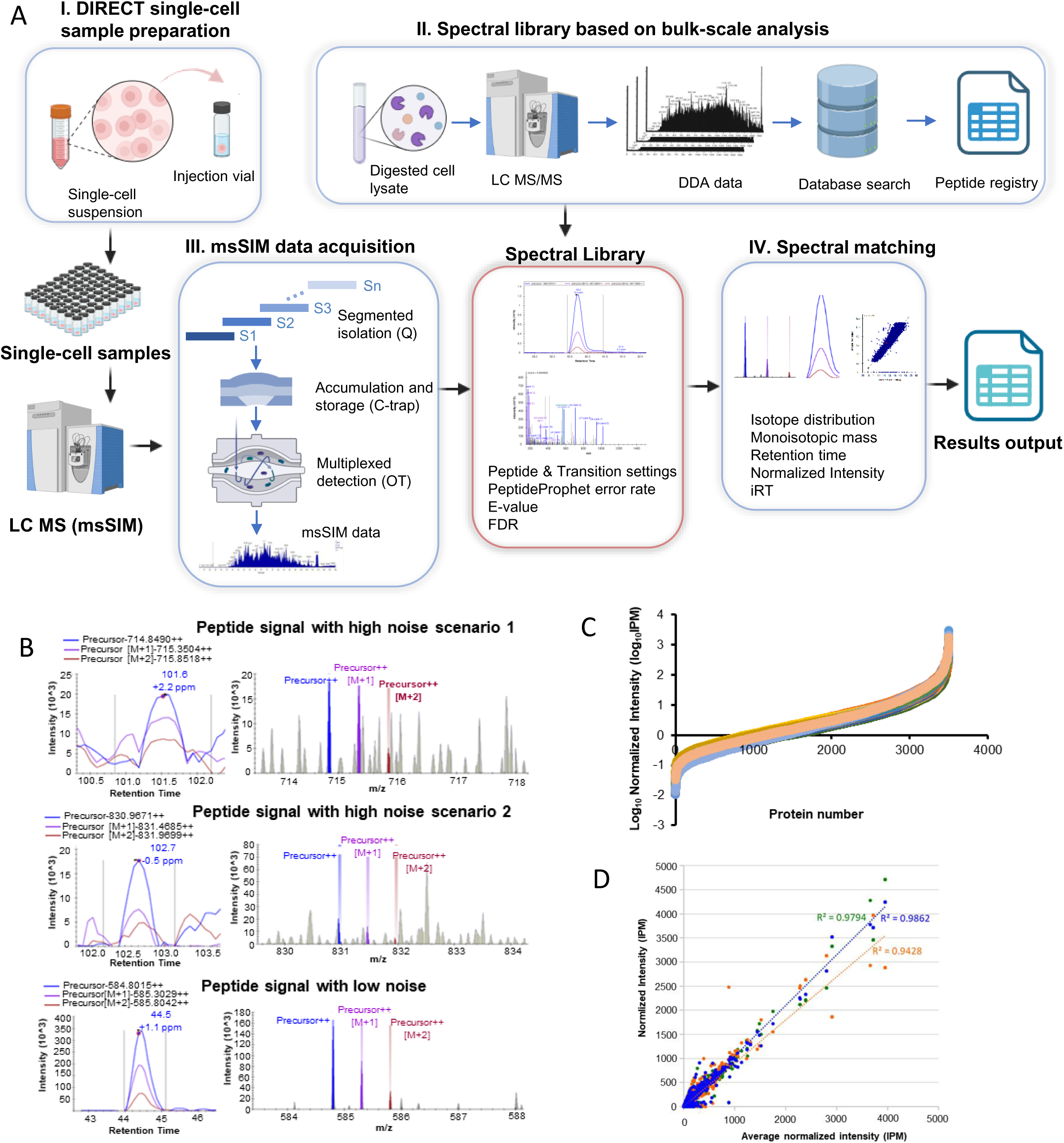
**The SLB-msSIM platform for highly sensitive and robust single-cell proteomic analysis. (**A) SLB-msSIM workflow. I. the “all-in-one” DIRECT method for single-cell preparation, II a high-resolution spectral library generated from a bulk-scale DDA analysis, III. LC-msSIM data acquisition, and IV. spectral library-based data analysis. (B) Examples of explicit peptide identification in the single-cell analysis of PANC-1 cells using SLB-msSIM. The left panels are peptide elution profiles and the right panels are peptide isotopic matchings. (C) The relative abundances of the common proteins identified in the individual HPDE cells. (D) Correlation of peptide intensities from three replicate analyses of a 500 pg Hela digest (equivalent to approximately 2-3 cells) using the SLB-msSIM.

For library construction, high-resolution DDA data acquired from the bulk-scale samples were used to establish a high-quality spectral library that serves as a registry for single-cell data analysis using spectral library matching. Single cells are dispensed, isolated, and processed with the DIRECT protocol, then analyzed with LC-msSIM with the parameters optimized for single-cell proteomic analysis. In the msSIM method, MS1 data of a single-cell sample was acquired by multiplexing segmented SIM acquisitions, in which precursor ions were consecutively selected by the quadrupole using a wide isolation window of 50 Th, accumulated in the C-trap, and then sent to the Orbitrap at once for single scan analysis.

The single-cell MS1 data obtained by the msSIM acquisition was matched against an empirically generated high-resolution spectral library for peptide identification based on monoisotopic mass, isotopic distribution, and retention time (hydrophobicity). These highly specific physicochemical characteristics of a peptide detected by chromatography and high-resolution mass spectrometry are well defined by the atomic composition and sequence of a peptide to permit highly accurate peptide matching.

### Spectral library matching aided by natural isotope distribution

It is important to note that in addition to an accurate mass and retention time (AMT), which may be sufficient for proteomic identification in a bulk-scale analysis[27, 28], the isotopic distribution of a peptide is a critical criterion for spectral library matching in a single-cell analysis, given the very low signal-to-noise ratio. Within a narrow window of AMT, an isotopic distribution matching can explicitly distinguish a unique peptide precursor signal (a fingerprint with a distinct distribution of isotopic peaks) from random noise signals (mostly single peaks) by quantitatively matching the well-defined isotopic envelope with the high-resolution spectral library[18]. **Figure 1B** exemplifies the explicit identification of peptides in both high-noise and low-noise environments in the single-cell analysis of PANC-1 cells using the SLB-msSIM. The peptide identification is critically aided by isotopic pattern matching, which can be quantified by Isotope Dot Product, to discriminate peptide signals from noise.

Studies have shown that multiple isolation windows over the full mass range can enhance the signal resolution and intensity, thus improve peptide identification[29]. A narrow range scan can significantly increase the sensitivity in MS1 acquisition compared to the wide range full-scan, especially on the detection of isotopic peaks[30]. Using the msSIM acquisition method, which multiplexes segmented SIMs to enhance precursor ion accumulation in the C-trap, dampen the noise level, and effectively separate confounding signals, we identified ∼3700 - ∼4300 proteins per single cell with stringent criteria in the analysis of nearly 200 individual cells with the SLB-msSIM. The proteins identified in single cells exceeded 5.5 orders of magnitude in protein abundance (**Figure 1C)**. Furthermore, by isolating a restricted m/z range, the selected ion monitoring mode can allow one to overcome the dynamic range limitations associated with trapping devices[31], and thereby enhance quantification. **Figure 1D** shows the correlations of peptide intensities from a replicate analysis of 500 picograms of HeLa digest (equivalent to the protein content of approximately 2-3 cells), demonstrating a high correlation and robustness in this pseudo single-cell analysis.

### Analysis of individual cancerous and normal pancreatic epithelial cells to interrogate functional differences in proteome landscape with single-cell resolution

We applied the SLB-msSIM method to analyze individual cells from four different pancreatic cancer lines (HPAF, AsPC-1, PANC-1, and MIA-PaCa2), along with normal pancreatic epithelial cells (HPDE). We applied highly stringent criteria for library construction and single cell proteomic identification. All peptides used for building the spectral library are filtered with a PeptideProphet error rate of ≤ 1% and an E-value of ≤ 0.05, resulting in FDRs ≤ 0.11% based on decoy database search. Using the “ID-Pair” approach[18, 32], we estimated that the false transfer rates (FTRs) from the libraries to the single-cell proteomes were between 1.55% – 16.18%. The average numbers of the peptides identified per protein for all cell lines are in the range of 4.7 – 6.5. **Figure 2A** summarizes the average numbers of peptides and proteins identified in each single cell for the five cell lines, ranging from 18228 to 25763 for peptide identification and 3697 to 4379 for protein identification. Using the HPDE cells as an example, the majority (>99%) of the peptides identified in the single cells have a mass deviation of less than 10 ppm from their theoretical values (**Figure 2B**).

**Figure 2.**
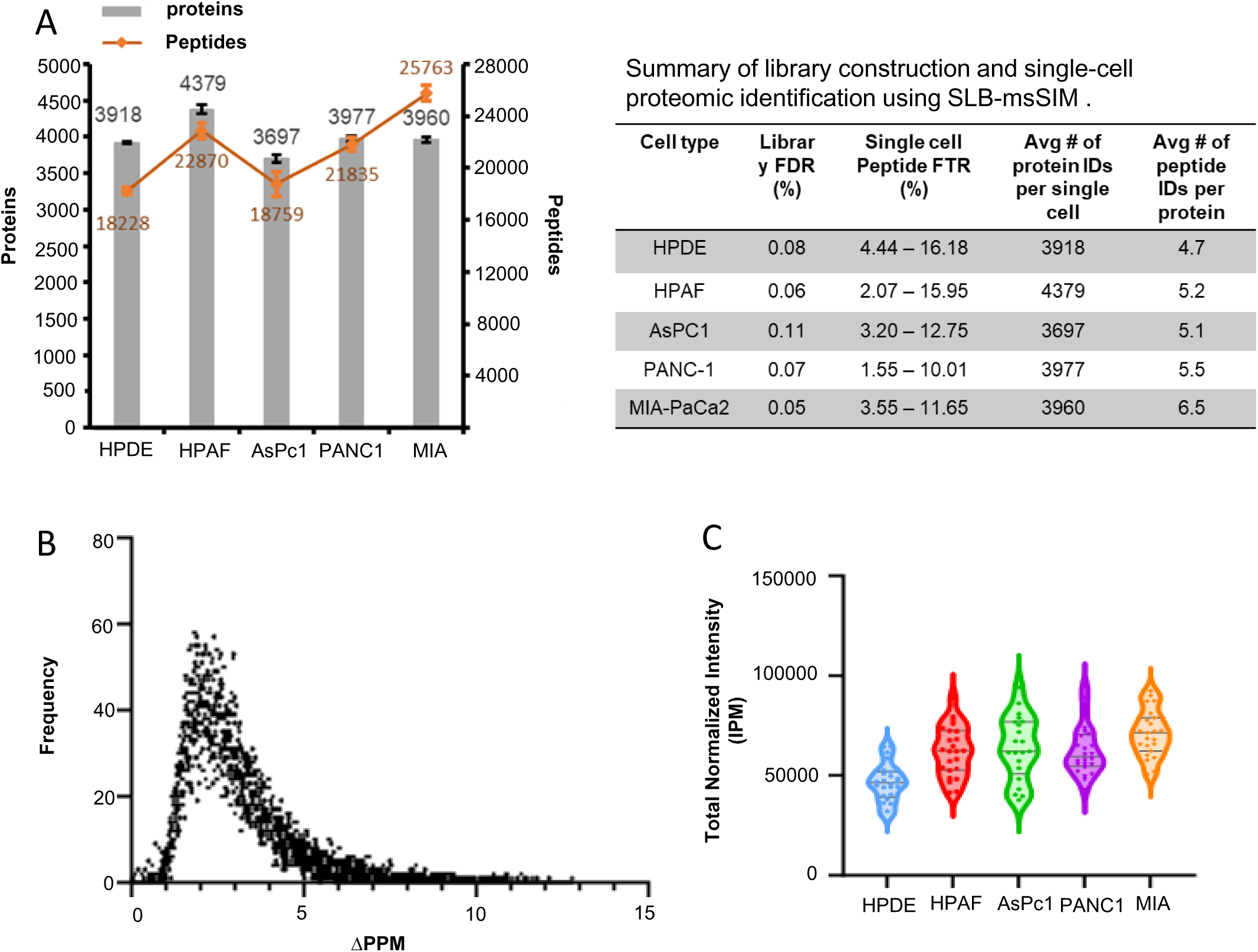
Single-cell proteomic profiling of individual cells from the HPDE, HPAF, AsPC-1, PANC-1, and MIA-PaCa2 cell lines. (A) Summary of library construction and single-cell proteomic identification using the SLB-msSIM. (B) The distribution of mass deviation (ΔPPM) of peptides identified in the HPDE single cells. (C) Comparison of normalized total protein contents in single cells from each cell line.

Notably, the comparison of the total protein content between the cancerous single cells and HPDE revealed that all four pancreatic cancer cell lines displayed more dispersed single-cell protein contents, while the protein contents in the HPDE cells were more homogeneous (**Figure 2C**). This observation is consistent with the heterogenous nature of these metastatic cancer cells[33–35], which could have more diverse gene expression in each cell. Furthermore, as a group, the protein contents of cancer cells appeared to be consistently higher than that of the normal HPDE cells (**Figure 2C**), possibly reflecting the altered proteomic landscape and reprogrammed biogenesis in these cancer cells.

Pairwise comparisons at single-cell level were conducted between each pancreatic cancer cell line (HPAF, AsPC-1, PANC-1, and MIA-PaCa2) and HPDE. The tNSE plots using the global protein profiles of individual cells distinctly exhibited well separations between normal and cancer cells for all four cancer cell lines (**Figure 3A, D, G, and J**). Proteins commonly identified between HPDE and each cancer cell line (**Figure 3B, E, H, and K**) underwent further analysis to identify up-regulated and down-regulated proteins. Volcano plots (**Figure 3C, F, I, and L**) demonstrate higher numbers of up-regulated proteins (fold change ≥ 2, p-value < 0.05) in all cancer cells compared to HPDE in comparison to the numbers of down-regulated proteins (fold change ≤ 0.5, p-value < 0.05). Many of these dysregulated protein expressions observed at the individual cell level are involved in metabolic pathways, DNA repair, protein synthesis, and other cellular processes[36–39]. It is also important to note that quantifications at single-cell level and bulk scale were not linearly correlated as we previously observed[18], highlighting the importance of single-cell proteomics as it may reflect different proteome landscapes that are not accessible from a bulk scale analysis.

**Figure 3.**
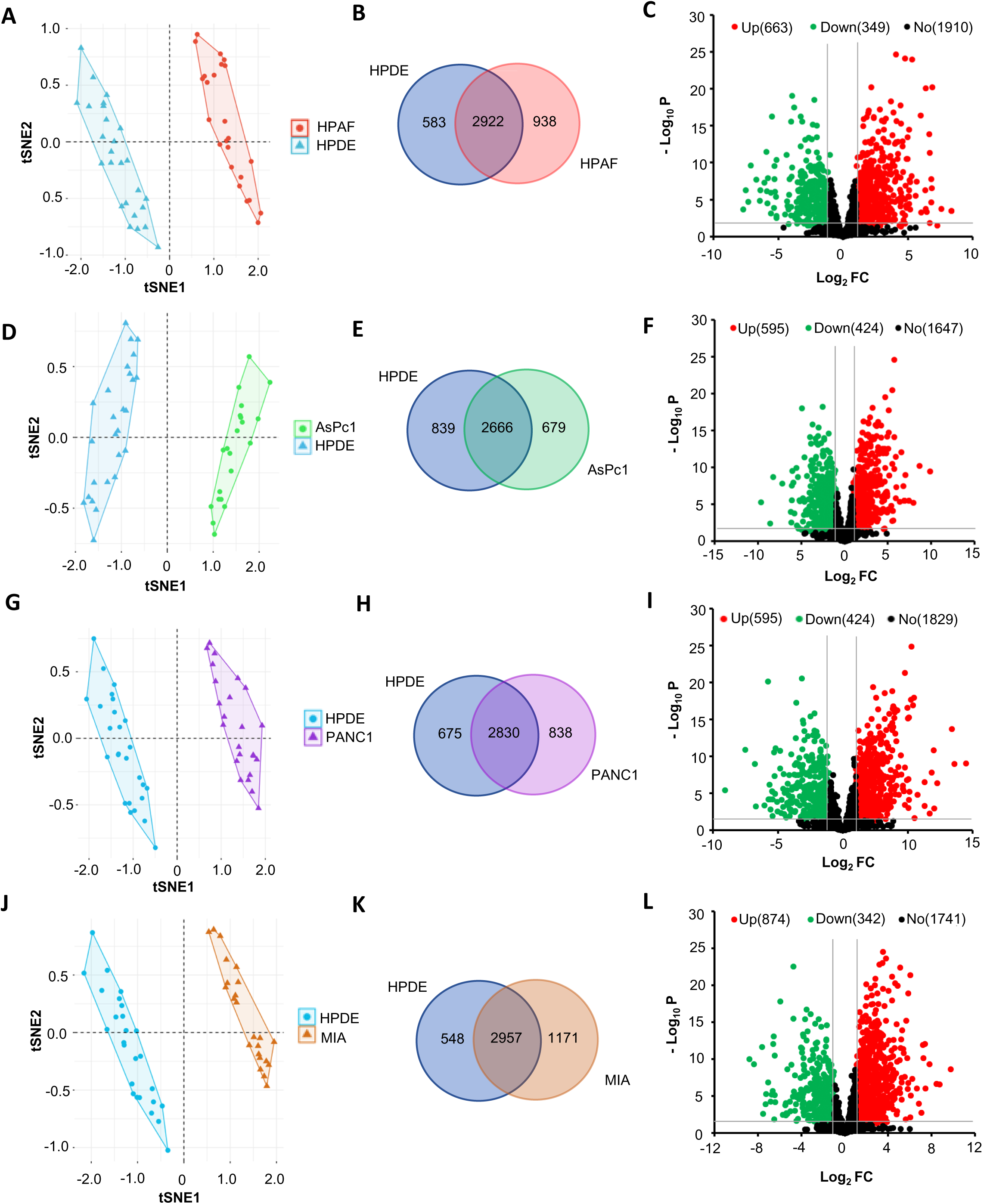
A pair-wise comparison of total protein profile of individual cells from each pancreatic cancer cell line (HPAF, AsPC-1, PANC-1, and MIA-PaCa2) with HPDE cells. t-SNE plots of (A) HPAF, (D) AsPc-1, (G) PANC1, and (J) MIA-PaCa2 compared to HPDE. Venn diagrams showing the common proteins identified in (B) HPAF, (E) AsPc-1, (H) PANC1, and (K) MIA-PaCa2 compared to HPDE. Volcano plots of (C) HPAF, (F) AsPc-1, (I) PANC1, and (L) MIA-PaCa2 compared to HPDE. Red and green dots indicate significantly upregulated and downregulated proteins based on their protein abundances, respectively (p-value < 0.05 and fold change ≥ 2).

Based on the GSEA analysis of the common 2300 proteins identified in the individual cancer cells from the four different pancreatic cancer cell lines and HPDE (**Figure 4A**), several cancer hallmarks were significantly (p-value < 0.05) enriched in the pancreatic cancer cells compared to the normal HPDE cells. These common cancer hallmarks include mitotic spindle, G2M checkpoint, fatty acid metabolism, MYC targets, and spermatogenesis (**Figure 4B**). The protein networks associated with the top enriched cancer hallmarks were exemplified in **Figure 4C**. It is not surprising that our data unveiled MYC as a hub gene in these four cancer cell lines as MYC is a master regulator of many cellular processes including proliferation, cell cycle[40] and metabolism[41]. It is downstream of KRAS signing[42], one of the most commonly activated pathways in pancreatic cancer[34, 43, 44]. Previous tissue proteomic studies in clinical samples have suggested that c-MYC is an important regulatory protein in the development of pancreatic intraepithelial neoplasia (PanIN) and pancreatic cancer[44, 45].

**Figure 4.**
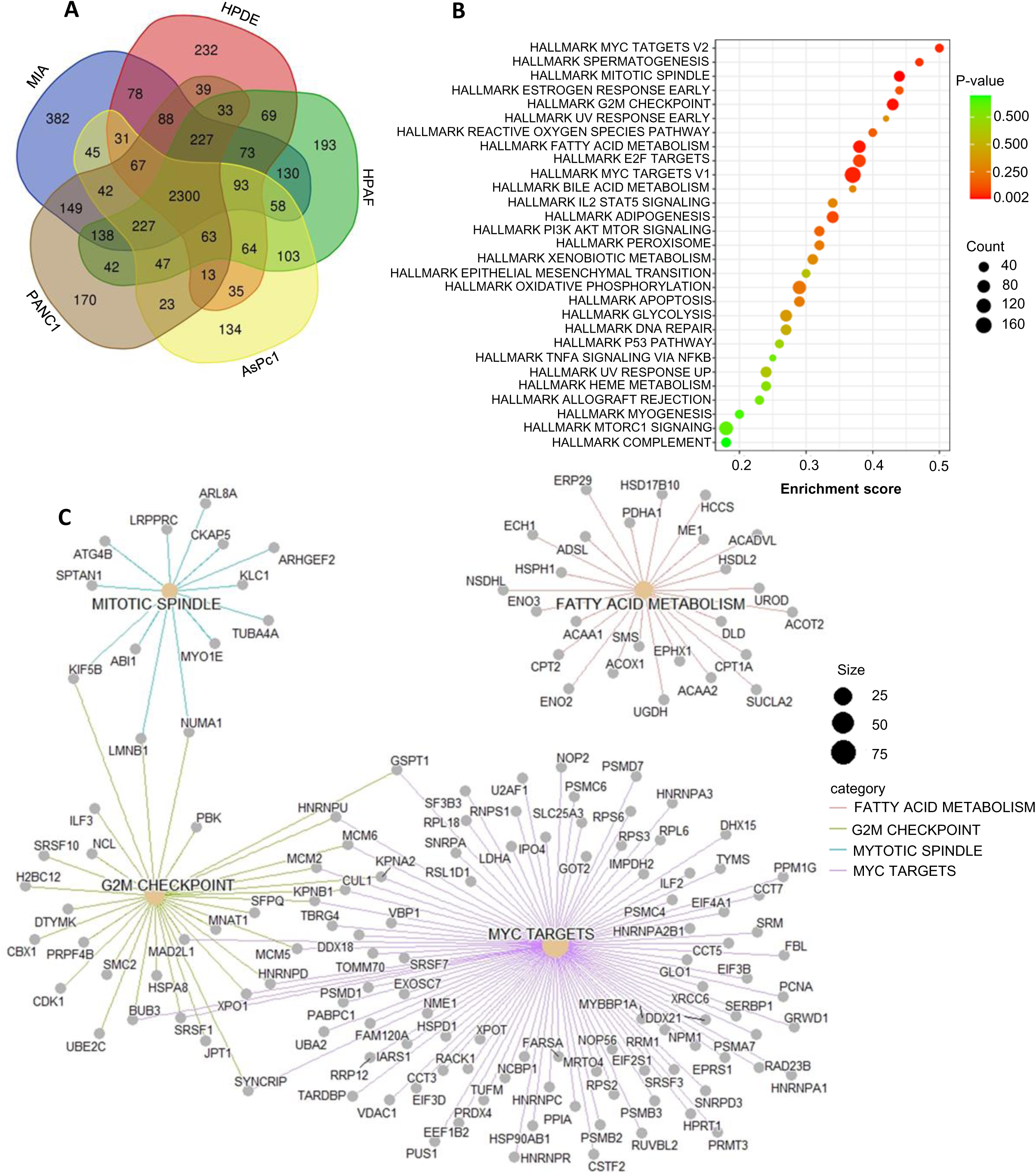
Enriched cancer hallmarks and associated protein networks revealed by functional analysis of the common proteins identified in cancer cells (HPAF, AsPC-1, PANC-1, and MIA-PaCa2) compared to normal HPDE cells. (A) Venn diagram to show the common proteins identified among the five cell lines. (B) A bubble plot showing cancer hallmark enrichment analysis of the overexpressed proteins identified through the GSEA analysis. (C) Protein networks of the top five hallmarks identified in the GSEA analysis based on the p-value. Proteins involved in MYC Targets V1 and MYC Targets V2 pathways were combined and represented together.

### Characterization of cellular heterogeneity in TGFβ1-induced EMT process and reversal

EMT is an important cancer hallmark as it is involved in cancer progression, invasion, metastasis, and drug resistance[19, 46, 47]. The reversal of EMT is aimed at mitigating the adverse effects of EMT in cancer and is considered an effective strategy to improve cancer treatment. Interrogation of single-cell trajectory and heterogeneity implicated in EMT and the reversal of EMT could give valuable insights into developing novel cancer therapies. In our EMT experiment, TGFβ1 stimulation was administered to PANC-1 cancer cells for 3 days for EMT induction; and subsequently, the TGFβ1 stimulation was discontinued to allow for reversal of EMT (**Figure 5A**). At the bulk scale, Western Blot analysis showed that the epithelial marker E-cadherin was downregulated, while the mesenchymal marker Snail was upregulated after TGFβ1 treatment for 3 days (D3) (**Figure 5B**). At day 7 (D7), four days after discontinued treatment of TGFβ1, the mesenchymal marker Snail returned to its basal expression levels similar to the control, while the epithelial marker E-cadherin remined to be downregulated (**Figure 5B**). The Western Blot data indicates a partial reversal of EMT at D7.

**Figure 5.**
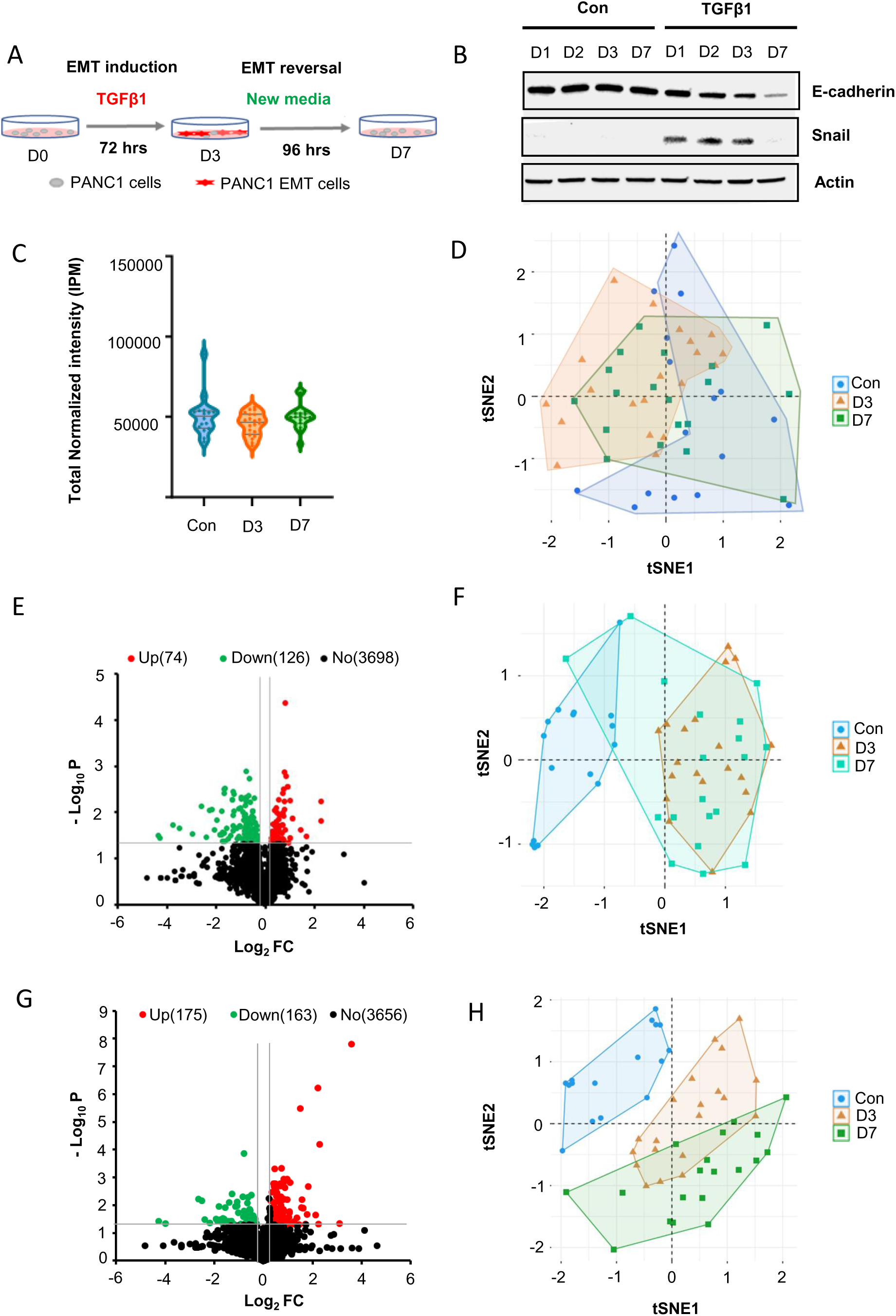
Changes of proteome in PANC-1 cells during the courses of EMT (TGFβ1 stimulation) and EMT reversal (removal of TGFβ1 stimulation). (A) The experimental setup. (B) The Western Blot indicating the bulk-scale protein expression for epithelial marker (E-cadherin) and mesenchymal marker (Snail) of PANC-1 cells during the courses of EMT and EMT reversal. Actin was used as the loading control. (C) Comparison of total normalized protein contents of individual cells in the EMT experiment. Con: untreated cells, D3: cells treated with TGFβ1 for 3 days and D7: cells after the removal of TGFβ1 for 4 days. (D) t-SNE plot based on total protein profiles of individual cells from Con, D3 and D7. (E) Volcano plot of protein abundances in the comparison of D3 vs. Con. (F) t-SNE plot based on the dysregulated proteins in D3 compared to Con. (G) Volcano plot of protein abundances in the comparison of D7 vs. Con. (H) t-SNE plot based on the dysregulated proteins in D7 compared to Con. Red and green dots in volcano plots indicate significantly upregulated and downregulated proteins based on their protein abundances, respectively (p-value < 0.05 and fold change ≥ 1. 2).

The SLB-msSIM method was applied to analyze the individual PANC-1 cells at Con (control with no TGFβ1 treatment), D3 (EMT) and D7 (EMT reversal). The single-cell proteomic data showed that the total proteome contents in the single cells of the PANC-1 cells changed during the course of EMT and EMT reversal (**Figure 5C**). The non-treated PANC-1 cancer cells (Con) were dispersed in a wider range compared to D3 (EMT) and D7 (EMT reversal) groups. The dispersed distribution of PANC-1 cells at basal level is consistent with the heterogeneous nature of cancer cells attributed to the altered proteome. The EMT induced by the TGFβ1 stimulation appeared to synchronize the total proteome contents of the PANC-1 cells after 3 days of treatment (D3). The removal of the stimulation for 4 days (D7) appeared to disturb this synchronization to a moderate extent, but it did not completely reverse the effect of EMT, as observed through the total protein content in single cells.

The t-SNE plots based on the global protein profiles of individual cells further showed that the majority of D3 (EMT) cells (orange triangles) were separated from the control cells (blue circles) (**Figure 5D**), indicating the distinct proteome alterations induced by TGFβ1 stimulation at the single cell level. After withdrawal of TGFβ1 treatment, individual cells at D7 (EMT reversal, green squares) exhibited a dispersed distribution that overlapped with both D3 (EMT) cells and control cells (**Figure 5D**), suggesting that while some of the D7 (EMT reversal) cells might have reverted to cellular states similar to those of the control cells, the majority of them might still retain proteomes similar to those of the D3 (EMT) cells. Furthermore, pairwise comparisons of proteome profiles were conducted to reveal the dysregulated proteins associated with EMT and its reversal (fold change ≥1.2, p-value < 0.05) in D3 (EMT) cells and D7 (EMT reversal) cells in comparison to the control cells **(Figure 5E and G**). The numbers of differential proteins associated with D3 (EMT) cells and D7 (EMT reversal) cells are 200 and 238, respectively. The t-SNE plots based on the dysregulated proteins associated with EMT cells confirmed that the majority of the D7 (EMT reversal) cells still retained the proteomes similar to those of the D3 (EMT) cells (**Figure 5F**). This observation is further supported by the t-SNE plots based on the dysregulated proteins associated with EMT reversal, which completely segregate the control cells from not only the D7 (EMT reversal) cells but also D3 (EMT) cells (**Figure 5H**).

The single-cell proteomic data indicated that the partial EMT reversal observed at the bulk-scale is likely primarily due to the heterogeneity in cellular trajectory. The individual single-cells stimulated with TGFβ1 and exhibiting an EMT status appeared to behave heterogeneously during the reversal of EMT. While the conditions and timings of TGFβ1 stimulation and its withdrawal may be crucial in reverting EMT markers and the associated proteome alterations to basal levels, the impacts of EMT on cellular proteome and functional homeostasis on individual cells may be multifactorial and heterogeneous.

## 4. DISCUSSION

We here report the development and application of the SLB-msSIM, a conceptually innovative MS1-based acquisition method with significantly enhanced sensitivity for single-cell analysis through spectra library matching. The msSIM method offers segmented, continuous data acquisition coverage over the entire scan range through multiplex ion accumulation, thus substantially enhancing the signal-to-noise ratio compared to full scan acquisition. The technology centers on the molecular phenomena where precursor ion signals are more intense than fragment ion signals generated through gas phase reactions in mass spectrometric analysis; and additionally, the distinct characteristics of a peptide precursor, including monoisotopic mass, isotopic distribution, and retention time (hydrophobicity), are uniquely associated with the peptide’s atomic composition and sequence. Notably, in addition to the accurate mass and retention time, the precise matching of the isotopic distribution of a peptide is a unique and critical criterion for distinguishing a peptide signal from random noise in single-cell analysis, given the very low signal-to-noise ratio. In the msSIM method, the precursor ions are sequentially isolated in multiple isolation cycles with the quadrupole using a wide isolation window and accumulated in the C-trap, where all ions stored are sent to the Orbitrap for detection in a single scan. Incorporation of the msSIM technique with the spectral library-based approach enabled the identification of ∼3700 - ∼4300 proteins per single cell with stringent criteria in the analysis of nearly 200 individual cells.

We have tested varying cell numbers (from 10 cells [1 ng] to 10,000 cells [1 µg]) for library construction using digested HeLa standards and observed a proportional increase in protein and peptide identifications with higher cell counts. We chose to use a bulk amount of 1 µg (approximately 10,000 cells) to generate the spectral library, primarily because it provides broader proteome coverage and strong, high-resolution isotopic signals that enhance spectral library matching for peptide identification.

We applied the SLB-msSIM method to two biological contexts: analyzing the individual cells from multiple pancreatic cancer cell lines in comparison to normal HPDE cells, and investigating the diverse cell trajectory during the EMT process and its reversal. Our single-cell proteomic data revealed the common and different functional characteristics among individual cells of PANC-1, MIA-PaCa2, AsPc-1 and HPAF pancreatic cancer cells in comparison to the normal HPDE. Furthermore, for the first time, we observed the cell trajectories of individual PANC-1 cells during EMT and its reversal, providing novel evidence of the impact of cellular heterogeneity on EMT reversal, a phenomenon relevant to cancer therapeutic development. The SLB-msSIM method offers a versatile alternative that significantly enhances the sensitivity and robustness of single-cell proteomic analysis and can be utilized across a wide range of instruments.

## 5. ASSOCIATED DATA

The mass spectrometric raw data have been deposited to the ProteomeXchange Consortium via PRIDE partner repository with the dataset identifier PXD055915.

## Acknowledgments

This work was supported in part by the Cancer Prevention & Research Institute of Texas (CPRIT) (RP210111), the National Institutes of Health (NIH) (R01CA180949 and R01CA276173), and the Rochelle and Max Levit endowment fund to SP. We thank Drs. Xiaodong Cheng and Fang Mei at University of Texas Health Science Center at Houston for kindly providing the HPDE cells.

## Supplementary Information

**Supplementary Table S1.** The iRT peptides used for internal calibration in the single-cell proteomic analysis.

## REFERENCES

[1] Liang, Y., Truong, T., Zhu, Y., Kelly, R. T., In-Depth Mass Spectrometry-Based Single-Cell and Nanoscale Proteomics. Methods Mol Biol 2021, 2185, 159–179.

[2] Li, Z. Y., Huang, M., Wang, X. K., Zhu, Y., et al., Nanoliter-Scale Oil-Air-Droplet Chip-Based Single Cell Proteomic Analysis. Anal Chem 2018, 90, 5430–5438.

[3] Williams, S. M., Liyu, A. V., Tsai, C. F., Moore, R. J., et al., Automated Coupling of Nanodroplet Sample Preparation with Liquid Chromatography-Mass Spectrometry for High-Throughput Single-Cell Proteomics. Anal Chem 2020, 92, 10588–10596.

[4] Kalxdorf, M., Muller, T., Stegle, O., Krijgsveld, J., IceR improves proteome coverage and data completeness in global and single-cell proteomics. Nat Commun 2021, 12, 4787.

[5] Gebreyesus, S. T., Siyal, A. A., Kitata, R. B., Chen, E. S., et al., Streamlined single-cell proteomics by an integrated microfluidic chip and data-independent acquisition mass spectrometry. Nat Commun 2022, 13, 37.

[6] Webber, K. G. I., Truong, T., Johnston, S. M., Zapata, S. E., et al., Label-Free Profiling of up to 200 Single-Cell Proteomes per Day Using a Dual-Column Nanoflow Liquid Chromatography Platform. Anal Chem 2022, 94, 6017–6025.

[7] Woo, J., Clair, G. C., Williams, S. M., Feng, S., et al., Three-dimensional feature matching improves coverage for single-cell proteomics based on ion mobility filtering. Cell Syst 2022, 13, 426–434 e424.

[8] Budnik, B., Levy, E., Harmange, G., Slavov, N., SCoPE-MS: mass spectrometry of single mammalian cells quantifies proteome heterogeneity during cell differentiation. Genome Biol 2018, 19, 161.

[9] Cheung, T. K., Lee, C. Y., Bayer, F. P., McCoy, A., et al., Defining the carrier proteome limit for single-cell proteomics. Nat Methods 2021, 18, 76–83.

[10] Campbell, A. J., Cakar, S., Palstrom, N. B., Riber, L. P., et al., A Carrier-Based Quantitative Proteomics Method Applied to Biomarker Discovery in Pericardial Fluid. Mol Cell Proteomics 2024, 23, 100812.

[11] Tsai, C. F., Zhao, R., Williams, S. M., Moore, R. J., et al., An Improved Boosting to Amplify Signal with Isobaric Labeling (iBASIL) Strategy for Precise Quantitative Single-cell Proteomics. Mol Cell Proteomics 2020, 19, 828–838.

[12] Schoof, E. M., Furtwangler, B., Uresin, N., Rapin, N., et al., Quantitative single-cell proteomics as a tool to characterize cellular hierarchies. Nat Commun 2021, 12, 3341.

[13] Cong, Y., Motamedchaboki, K., Misal, S. A., Liang, Y., et al., Ultrasensitive single-cell proteomics workflow identifies >1000 protein groups per mammalian cell. Chem Sci 2020, 12, 1001–1006.

[14] Brunner, A. D., Thielert, M., Vasilopoulou, C., Ammar, C., et al., Ultra-high sensitivity mass spectrometry quantifies single-cell proteome changes upon perturbation. Mol Syst Biol 2022, 18, e10798.

[15] Ctortecka, C., Clark, N. M., Boyle, B. W., Seth, A., et al., Automated single-cell proteomics providing sufficient proteome depth to study complex biology beyond cell type classifications. Nat Commun 2024, 15, 5707.

[16] Wang, J., Tan, H., Fu, Y., Mishra, A., et al., Evaluation of Protein Identification and Quantification by the diaPASEF Method on timsTOF SCP. J Am Soc Mass Spectrom 2024, 35, 1253–1260.

[17] Dwivedi, P., Rose, C. M., Understanding the effect of carrier proteomes in single cell proteomic studies-key lessons. Expert Rev Proteomics 2022, 19, 5–15.

[18] Senavirathna, L., Ma, C., Chen, R., Pan, S., Spectral Library-Based Single-Cell Proteomics Resolves Cellular Heterogeneity. Cells 2022, 11.

[19] Shibue, T., Weinberg, R. A., EMT, CSCs, and drug resistance: the mechanistic link and clinical implications. Nat Rev Clin Oncol 2017, 14, 611–629.

[20] Nakasuka, F., Tabata, S., Sakamoto, T., Hirayama, A., et al., TGF-beta-dependent reprogramming of amino acid metabolism induces epithelial-mesenchymal transition in non-small cell lung cancers. Commun Biol 2021, 4, 782.

[21] Eng, J. K., Jahan, T. A., Hoopmann, M. R., Comet: an open-source MS/MS sequence database search tool. Proteomics 2013, 13, 22–24.

[22] Deutsch, E. W., Mendoza, L., Shteynberg, D. D., Hoopmann, M. R., et al., Trans-Proteomic Pipeline: Robust Mass Spectrometry-Based Proteomics Data Analysis Suite. J Proteome Res 2023, 22, 615–624.

[23] Nesvizhskii, A. I., Protein identification by tandem mass spectrometry and sequence database searching. Methods Mol. Biol 2007, 367, 87–119.

[24] Maclean, B., Tomazela, D. M., Shulman, N., Chambers, M., et al., Skyline: an open source document editor for creating and analyzing targeted proteomics experiments. Bioinformatics 2010, 26, 966–968.

[25] Parker, S. J., Rost, H., Rosenberger, G., Collins, B. C., et al., Identification of a Set of Conserved Eukaryotic Internal Retention Time Standards for Data-independent Acquisition Mass Spectrometry. Mol Cell Proteomics 2015, 14, 2800–2813.

[26] Subramanian, A., Tamayo, P., Mootha, V. K., Mukherjee, S., et al., Gene set enrichment analysis: a knowledge-based approach for interpreting genome-wide expression profiles. Proc Natl Acad Sci U S A 2005, 102, 15545–15550.

[27] May, D., Liu, Y., Law, W., Fitzgibbon, M., et al., Peptide sequence confidence in accurate mass and time analysis and its use in complex proteomics experiments. J Proteome Res 2008, 7, 5148–5156.

[28] Zimmer, J. S., Monroe, M. E., Qian, W. J., Smith, R. D., Advances in proteomics data analysis and display using an accurate mass and time tag approach. Mass Spectrom Rev 2006, 25, 450–482.

[29] Egertson, J. D., Kuehn, A., Merrihew, G. E., Bateman, N. W., et al., Multiplexed MS/MS for improved data-independent acquisition. Nat. Methods 2013, 10, 744–746.

[30] Kaufmann, A., Analytical performance of the various acquisition modes in Orbitrap MS and MS/MS. J Mass Spectrom 2018, 53, 725–738.

[31] Gallien, S., Duriez, E., Crone, C., Kellmann, M., et al., Targeted proteomic quantification on quadrupole-orbitrap mass spectrometer. Mol Cell Proteomics 2012, 11, 1709–1723.

[32] Lim, M. Y., Paulo, J. A., Gygi, S. P., Evaluating False Transfer Rates from the Match-between-Runs Algorithm with a Two-Proteome Model. J Proteome Res 2019, 18, 4020–4026.

[33] Pasha, N., Turner, N. C., Understanding and overcoming tumor heterogeneity in metastatic breast cancer treatment. Nat Cancer 2021, 2, 680–692.

[34] Deer, E. L., Gonzalez-Hernandez, J., Coursen, J. D., Shea, J. E., et al., Phenotype and genotype of pancreatic cancer cell lines. Pancreas 2010, 39, 425–435.

[35] Sasaki, N., Gomi, F., Hasegawa, F., Hirano, K., et al., Characterization of the metastatic potential of the floating cell component of MIA PaCa-2, a human pancreatic cancer cell line. Biochem Biophys Res Commun 2020, 522, 881–888.

[36] Pessoa, J., Martins, M., Casimiro, S., Perez-Plasencia, C., Shoshan-Barmatz, V., Editorial: Altered Expression of Proteins in Cancer: Function and Potential Therapeutic Targets. Front Oncol 2022, 12, 949139.

[37] Hanahan, D., Weinberg, R. A., The hallmarks of cancer. Cell 2000, 100, 57–70.

[38] Bradner, J. E., Hnisz, D., Young, R. A., Transcriptional Addiction in Cancer. Cell 2017, 168, 629–643.

[39] DeBerardinis, R. J., Lum, J. J., Hatzivassiliou, G., Thompson, C. B., The biology of cancer: metabolic reprogramming fuels cell growth and proliferation. Cell Metab 2008, 7, 11–20.

[40] Garcia-Gutierrez, L., Delgado, M. D., Leon, J., MYC Oncogene Contributions to Release of Cell Cycle Brakes. Genes (Basel) 2019, 10.

[41] Miller, D. M., Thomas, S. D., Islam, A., Muench, D., Sedoris, K., c-Myc and cancer metabolism. Clin Cancer Res 2012, 18, 5546–5553.

[42] Ala, M., Target c-Myc to treat pancreatic cancer. Cancer Biol Ther 2022, 23, 34–50.

[43] Zhang, Z., Zhang, H., Liao, X., Tsai, H. I., KRAS mutation: The booster of pancreatic ductal adenocarcinoma transformation and progression. Front Cell Dev Biol 2023, 11, 1147676.

[44] Pan, S., Brentnall, T. A., Chen, R., Proteome alterations in pancreatic ductal adenocarcinoma. Cancer Lett 2020, 469, 429–436.

[45] Pan, S., Chen, R., Reimel, B. A., Crispin, D. A., et al., Quantitative proteomics investigation of pancreatic intraepithelial neoplasia. Electrophoresis 2009, 30, 1132–1144.

[46] Williams, E. D., Gao, D., Redfern, A., Thompson, E. W., Controversies around epithelial-mesenchymal plasticity in cancer metastasis. Nat Rev Cancer 2019, 19, 716–732.

[47] De Las Rivas, J., Brozovic, A., Izraely, S., Casas-Pais, A., et al., Cancer drug resistance induced by EMT: novel therapeutic strategies. Arch Toxicol 2021, 95, 2279–2297.

