## Supplemental Materials for "SLB-msSIM: a spectral library-based multiplex segmented SIM platform for single-cell proteomic analysis"

### Supplementary Information

**Supplementary Table 1.** The iRT peptides used for internal calibration

| |
|---|
| AGFAGDDAPR |
| ATAGDTHLGGEDFDNR |
| LVLVGDGGTGK |
| VAVVAGYGDVGK |
| IGGIGTVPVGR |
| AVFPSIVGRPR |
| TTPSYVAFTDTER |
| IGLFGGAGVGK |
| VCENIPIVLCGNK |
| DAGTIAGLNVLR |
| SYELPDGQVITIGNER |
